# NanoSpring: reference-free lossless compression of nanopore sequencing reads using an approximate assembly approach

**DOI:** 10.1101/2021.06.09.447198

**Authors:** Qingxi Meng, Shubham Chandak, Yifan Zhu, Tsachy Weissman

## Abstract

The amount of data produced by genome sequencing experiments has been growing rapidly over the past several years, making compression important for efficient storage, transfer and analysis of the data. In recent years, nanopore sequencing technologies have seen increasing adoption since they are portable, real-time and provide long reads. However, there has been limited progress on compression of nanopore sequencing reads obtained in FASTQ files since most existing tools are either general-purpose or specialized for short read data.

We present NanoSpring, a reference-free compressor for nanopore sequencing reads, relying on an approximate assembly approach. We evaluate NanoSpring on a variety of datasets including bacterial, metagenomic, plant, animal, and human whole genome data. For recently basecalled high quality nanopore datasets, NanoSpring, which focuses only on the base sequences in the FASTQ file, uses just 0.35-0.65 bits per base which is 3-6x lower than general purpose compressors like gzip. NanoSpring is competitive in compression ratio and compression resource usage with the state-of-the-art tool CoLoRd while being significantly faster at decompression. NanoSpring is available on GitHub at https://github.com/qm2/NanoSpring.

## Introduction

The rapid decrease in the cost of genome sequencing has led to an explosion in the amount of data produced by these experiments, with the raw data usually requiring the most space for storage. The raw sequencing data is obtained in the form of reads with sequencing depth/coverage often being 30x or higher. A typical human whole genome sequencing experiment can produce 100s of GBs of data in FASTQ files. Given the high sequencing depth, there is much redundancy to be exploited in the reads, and several specialized compressors like SPRING^1^ and PgRC^2^ have been developed for this data. The typical approach used by these compressors is to efficiently build an approximate assembly using the reads and then store this assembly along with the encoding of the reads with respect to the assembly.

While the existing compressors have been mostly built for short-read sequencers such as Illumina, in recent years, nanopore sequencing, specifically using Oxford Nanopore Technologies (ONT) sequencers^3^, has seen increasing adoption since it is portable, real-time and provides long reads. However, there has been limited progress on compression of nanopore sequencing reads. Most existing works like SPRING and PgRC operate under the assumption that the reads are short (~100s of bases) and low-error (with most errors being substitutions). On the other hand, nanopore reads are much longer (often over hundreds of thousands of bases long), and have a much higher error rate, including substitution, insertion, and deletion errors from the basecalling process that converts the raw current signal to the read sequences^4^. However, the error rate has fallen dramatically in the recent years with the advent of deep learning based basecallers which achieve median error rate close to 5% or better^5^, suggesting that a similar approximate assembly approach with some adaptations can be applied to nanopore sequencing reads.

There have been a few works on compression of nanopore FASTQ data in recent years. ENANO^6^ focuses mostly on quality score compression, and uses a context-based model followed by arithmetic coding for read sequences. Note that while quality scores occupy a significant amount of space even after compression, we focus on read sequences due to the relative lack of research in this area, and since quality scores are often ignored by downstream tools like minimap2^7^. Quality scores have also been compressed lossily without an impact on the downstream performance for short-read technologies^8,9^ and more recently for nanopore itself^10,11^. RENANO^12^ is a recent reference-based compressor that achieves significantly better compression for read sequences, but is limited to aligned data with a reference available. Most recently, CoLoRd^10^ included both a reference-free and reference-based compressor using overlap graph based approach, achieving significant improvement over ENANO in the reference-free mode at the cost of higher resource usage. In addition, CoLoRd also includes analysis of lossy quality value compression showcasing that the impact on downstream performance is minimal.

In this work, we present NanoSpring, which is a lossless reference-free compressor for nanopore sequencing reads. NanoSpring uses an approximate assembly approach partly inspired by existing assembly algorithms but adapted for significantly better performance, especially for the recent higher quality datasets. On recent human whole genome datasets, NanoSpring achieves close to 3-6x improvement in compression as compared to general purpose compressors and ENANO. NanoSpring is competitive in compression ratio and compression resource usage with the state-of-the-art tool CoLoRd while being significantly faster at decompression. NanoSpring is available as an open-source tool on GitHub, requires only a FASTQ file as input for compression, and does not compress read identifiers or quality values.

## Results and discussion

### NanoSpring algorithm overview

NanoSpring is a lossless compressor for nanopore read sequences and does not require an external reference for compression. NanoSpring relies on an approximate-assembly approach, where we first assemble the reads into contigs, obtain the consensus sequence for each contig, and finally store the consensus sequence and encode the reads with respect to the consensus sequence. Parts of the algorithm were inspired by the MinHash-based assembler MHAP^13^, with suitable adjustments to the parameters to achieve orders of magnitude speedup over the assembler while still obtaining a sufficiently accurate assembly for compression purposes.

NanoSpring first converts the reads into an efficient 2 bits/base representation on disk (ignoring read identifiers and quality values in the FASTQ file). Next, NanoSpring indexes the reads using MinHash which enables efficient lookup of reads overlapping a given sequence, effectively handling substitution, insertion, and deletion errors. Once the index is constructed, NanoSpring attempts to build contigs consisting of overlapping reads. The contigs are represented using consensus graphs with each read corresponding to a path on the graph. The contig is built by greedily searching the MinHash index for reads that overlap with the current consensus sequence of the graph, and adding the candidate reads to the graph using minimap7^7^ alignment. Finally, the consensus sequence and the errors in the reads with respect to the consensus sequence are written to separate streams and compressed using general-purpose compressors. This process is illustrated in Figure 1a. Any reads that remain left out of this process (*lone reads*) are simply encoded directly. The decompression process is quite simple: the decompressor first obtains the consensus sequence and error streams using the general-purpose decompressor. Then it applies the errors to the appropriate parts of the consensus sequence to obtain the reads, using disk based sorting to recover the reads in the original ordering while minimizing memory usage. More detailed description of the various stages, design choices and parameters is available in *Methods*.

**Figure 1.**
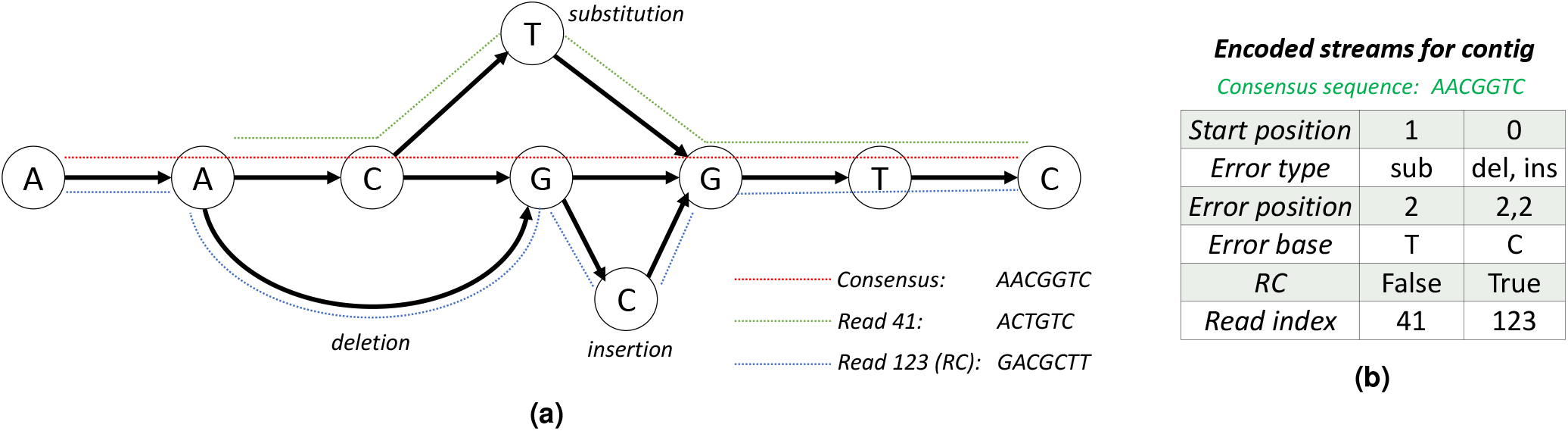
(a) Consensus graph for a contig showing the consensus sequence (path shown in red) and two reads. The read with the path shown in green has index 41 in the FASTQ file, and has a substitution compared to the consensus sequence. The read with the path shown in blue has index 123 in the FASTQ file, is reverse complemented with respect to the consensus sequence, and has a deletion and an insertion. (b) The encoding of the contig into multiple streams. Note that the error position is 0-indexed and delta coded, and the error base needs to be stored only for insertions and substitutions.

### Experimental setup

We tested NanoSpring on several real datasets that cover a variety of organisms with different genome lengths sequenced at varying depths of coverage. We compare NanoSpring to the current state-of-the-art reference-free compressors for nanopore FASTQ files, ENANO^6^ and CoLoRd^10^ and to pigz (https://zlib.net/pigz/) which is a multithreaded version of the general-purpose compressor Gzip. While ENANO compresses the entire FASTQ file including the read sequences, quality values and read identifiers, we only focus on the compressed size for the read sequences. We note that ENANO supports multiple compression levels, and we use the default one since it is significantly faster than the maximum compression level with minimal difference in read sequence compression (< 1%). We ran reference-free CoLoRd in the balanced mode with quality and read identifier compression mode set to none. All experiments were run on an Ubuntu 18.04.5 server with 40 Intel Xeon processors (2.2 GHz) and 260 GB RAM. The tools were run with 20 threads unless specified otherwise. Details on installing and running the various tools are provided in Supplementary data.

#### Datasets

The datasets used for experiments are listed in Table 1. These include bacterial, metagenomic, animal, plant, and human datasets. Further details on obtaining these datasets are provided in the Supplementary data. We included some standard datasets including the NA12878 dataset (*hs1*) and the Zymo microbial standard (*zymo*), but we largely focused on datasets basecalled with more recent tools that provided much higher quality and better scope for compression, including a R10.4 dataset (*hs4*). To further test the impact of the basecaller on the compression rate, we basecalled the *S. aureus* dataset using three modes and compare the results later. We also looked into the impact of coverage on the compression ratio for the *M. acuminata (banana)* and CHM13 dataset (*hs3*) which were available at a high initial coverage (results shown later)

**Table 1.** Datasets used for experiments. The uncompressed size refers to the file size obtained by removing the quality scores and sequence identifiers from the FASTQ files. N50 is a robust measure for read lengths, with the reads with length above the N50 metric capturing 50% of the data. The *zymo* dataset was obtained from a bonito basecalling of the Zymo sample (details at https://github.com/Kirk3gaard/2020-05-20_ZymoMock_Q20EA). The *hs2* and *hs4* datasets were was obtained from the ONT Open Datasets (https://labs.epi2me.io/gm24385_2020.09 / and https://labs.epi2me.io/gm24385_q20_2021.10/, respectively).

| Dataset Name | Species | Sample | Genome Size (Mbp) | Coverage | Number of Reads (M) | Average Read Length (Kb) | N50 (Kb) | Uncompressed Size (GB) | Source |
| --- | --- | --- | --- | --- | --- | --- | --- | --- | --- |
| *sa* | *S. aureus* | CAS28_02 | 2.9 | 84x | 0.01 | 21.99 | 24.8 | 0.24 | <sup>4</sup> |
| *zymo* | Metagenomic | SAMEA8172073 | - | - | 5.44 | 3.30 | 4.9 | 18.0 | See caption |
| *snail* | *C. squamiferum* | SAMN10963494 | 356 | 139x | 7.45 | 6.65 | 7.3 | 49.5 | <sup>14</sup> |
| *banana* | *M. acuminata* | SAMEA6104609 | 523 | 177x | 5.19 | 17.9 | 31.6 | 92.7 | <sup>15</sup> |
| *hs1* | *H. sapiens* | NA12878 | 3,200 | 42x | 15.7 | 8.48 | 13.6 | 132.9 | <sup>3</sup> |
| *hs2* | *H. sapiens* | GM24385 | 3,200 | 26x | 3.44 | 24.5 | 46.5 | 84.2 | See caption |
| *hs3* | *H. sapiens* | CHM13 | 3,200 | 23x | 5.90 | 12.6 | 58.7 | 74.2 | <sup>16</sup> |
| *hs4* | *H. sapiens* | GM24385 | 3,200 | 40x | 9.6 | 12.3 | 22.4 | 117.6 | See caption |

#### Compression results

Table 2 shows the compression results for Gzip, ENANO, CoLoRd, and NanoSpring on the datasets. We observe that Gzip and ENANO perform consistently across the datasets, achieving around 2.2 and 1.9 bits/base, respectively (with ~2 bits/base being achievable with a fixed-length encoding). NanoSpring provides much better compression, getting below 0.7 bits/base for the human datasets (*hs2, hs3*) and to 0.35 bits/base for *hs4*, which is around 3-6x better than Gzip and ENANO. Although CoLoRd generally provides better compression results than NanoSpring, NanoSpring is competitive for the recent human datasets with low error rates (e.g. *hs2, hs3, hs4*). In absolute terms, NanoSpring compresses the 84 GB *hs2* dataset to less than 7 GB. For most other datasets, NanoSpring achieves close to 2x improvement over the Gzip and Enano. Note that the compression results for the *hs1* dataset are significantly worse, although NanoSpring still outperforms Gzip and ENANO. This can be explained by the fact that this dataset was obtained using an older basecaller with appreciably higher error rates. Given the steady improvement in basecaller quality over the years^4^, we can expect the performance of NanoSpring to improve further in the future (the impact of basecaller error rate on NanoSpring performance is discussed below).

**Table 2.** Compression results for read sequences using Gzip, ENANO and NanoSpring.

| Dataset Name | Coverage | Uncompressed size<br>(GB) | Compressed size in bits/base |  |  |  |
| --- | --- | --- | --- | --- | --- | --- |
|  |  |  | Gzip | ENANO | CoLoRd | NanoSpring |
| *sa* | 84x | 0.24 | 2.29 | 1.89 | 0.48 | 0.49 |
| *zymo* | - | 18.0 | 2.33 | 1.96 | 0.49 | 0.60 |
| *snail* | 139x | 49.5 | 2.14 | 1.80 | 0.95 | 1.06 |
| *banana* | 177x | 92.7 | 2.28 | 1.93 | 0.91 | 1.08 |
| *hs1* | 42x | 132.9 | 2.24 | 1.89 | 1.13 | 1.47 |
| *hs2* | 26x | 84.2 | 2.20 | 1.87 | 0.60 | 0.67 |
| *hs3* | 23x | 74.2 | 2.18 | 1.86 | 0.59 | 0.68 |
| *hs4* | 40x | 117.6 | 2.18 | 1.86 | 0.33 | 0.35 |

#### Time and memory usage

Table 3 shows the time and peak memory usage for the compression and decompression using the four tools, all running on 20 threads. We note that ENANO does not provide a mode for compressing only the read sequences, and so the time and memory usage numbers include the compression of quality scores and read identifiers. Despite this, we see that ENANO significantly outperforms CoLoRd and NanoSpring in terms of compression time/memory and decompression memory. Gzip also uses much less time and memory than CoLoRd and NanoSpring. Among the approximate assembly based compressors, we observe that CoLoRd generally does a bit better than NanoSpring on compression time and memory usage. On the other hand, NanoSpring is faster and more memory efficient during decompression as compared to CoLoRd, requiring around 11 minutes and 5.6 GB peak memory for a 26x human dataset (*hs2*).

**Table 3.**
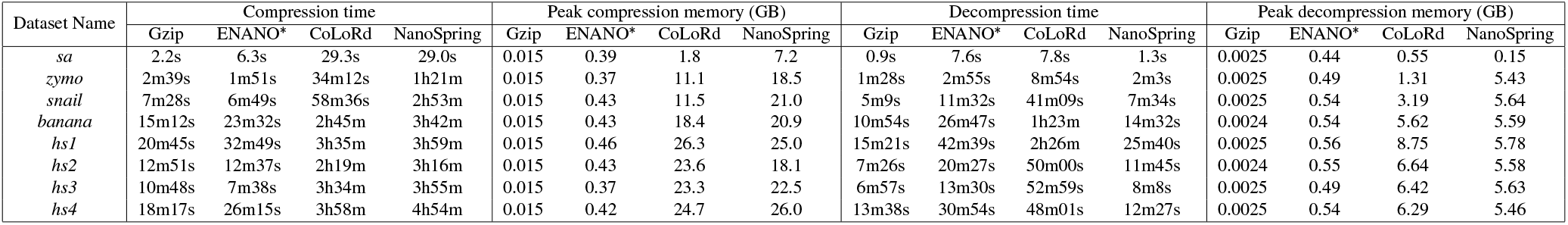
Time and peak memory requirements for compression and decompression. *ENANO figures include the time/memory usage to compress/decompress the entire FASTQ file including read sequences, quality values and read identifiers.

| Dataset Name | Compression time |  |  |  | Peak compression memory (GB) |  |  |  | Decompression time |  |  |  | Peak decompression memory (GB) |  |  |  |
| --- | --- | --- | --- | --- | --- | --- | --- | --- | --- | --- | --- | --- | --- | --- | --- | --- |
|  | Gzip | ENANO* | CoLoRd | NanoSpring | Gzip | ENANO* | CoLoRd | NanoSpring | Gzip | ENANO* | CoLoRd | NanoSpring | Gzip | ENANO* | CoLoRd | NanoSpring |
| *sa* | 2.2s | 6.3s | 29.3s | 29.0s | 0.015 | 0.39 | 1.8 | 7.2 | 0.9s | 7.6s | 7.8s | 1.3s | 0.0025 | 0.44 | 0.55 | 0.15 |
| *zymo* | 2m39s | 1m51s | 34m12s | 1h21m | 0.015 | 0.37 | 11.1 | 18.5 | 1m28s | 2m55s | 8m54s | 2m3s | 0.0025 | 0.49 | 1.31 | 5.43 |
| *snail* | 7m28s | 6m49s | 58m36s | 2h53m | 0.015 | 0.43 | 11.5 | 21.0 | 5m9s | 11m32s | 41m09s | 7m34s | 0.0025 | 0.54 | 3.19 | 5.64 |
| *banana* | 15m12s | 23m32s | 2h45m | 3h42m | 0.015 | 0.43 | 18.4 | 20.9 | 10m54s | 26m47s | 1h23m | 14m32s | 0.0024 | 0.54 | 5.62 | 5.59 |
| *hs1* | 20m45s | 32m49s | 3h35m | 3h59m | 0.015 | 0.46 | 26.3 | 25.0 | 15m21s | 42m39s | 2h26m | 25m40s | 0.0025 | 0.56 | 8.75 | 5.78 |
| *hs2* | 12m51s | 12m37s | 2h19m | 3h16m | 0.015 | 0.43 | 23.6 | 18.1 | 7m26s | 20m27s | 50m00s | 11m45s | 0.0024 | 0.55 | 6.64 | 5.58 |
| *hs3* | 10m48s | 7m38s | 3h34m | 3h55m | 0.015 | 0.37 | 23.3 | 22.5 | 6m57s | 13m30s | 52m59s | 8m8s | 0.0025 | 0.49 | 6.42 | 5.63 |
| *hs4* | 18m17s | 26m15s | 3h58m | 4h54m | 0.015 | 0.42 | 24.7 | 26.0 | 13m38s | 30m54s | 48m01s | 12m27s | 0.0025 | 0.54 | 6.29 | 5.46 |

The high resource usage for NanoSpring during compression is due to the approximate assembly process which provides the gains in compression, and is on a similar scale as previous works following this approach for short read^s1,2^. The memory usage consists of the MinHash index and the consensus graph. We typically observed that the time and memory usage scaled linearly with the dataset size, although there is some variability in the contig generation stage. For the 26x human dataset *hs2*, NanoSpring requires ~3 hours (~70 CPU hours) and 18.1 GB memory, which is an order of magnitude smaller than the requirements for genome assembly. For example, wtdbg2^17^, a recent efficient assembler requires over 1000 CPU hours and 200 GB memory for nanopore human datasets with ~35x coverage.

#### Contribution of streams to compressed size

Figure 2 shows the contribution of the various streams to the total compressed size for the datasets from Table 1. We focus on the consensus sequence, the error streams (position, type, erroneous base) and the lone reads (i.e., reads for which no matches were found). The remaining streams contribute less than 1% to the total size and are omitted here for clarity. We first note that the error streams take up close to 0.5 bits/base for most datasets, while the contribution of the consensus sequence is smaller. The contribution of the lone read stream varies a lot between datasets and is quite high for the *snail, banana* and *hs1* datasets where NanoSpring has relatively worse compression performance. We believe that this is associated with the data quality since we see a drastic reduction in the lone reads for datasets basecalled with the latest basecallers (*sa*, *hs2*, *hs3* and *hs4*). For the high-quality *hs4* dataset where we get out best performance, the contribution of lone reads is negligible. Finally, we note that datasets with higher coverage have a smaller contribution from the consensus sequence (e.g., the *zymo* dataset which has extremely high average coverage across the bacterial species). This is expected theoretically since the genome size is a fixed constant while the error streams grow linearly as we get more reads.

**Figure 2.**
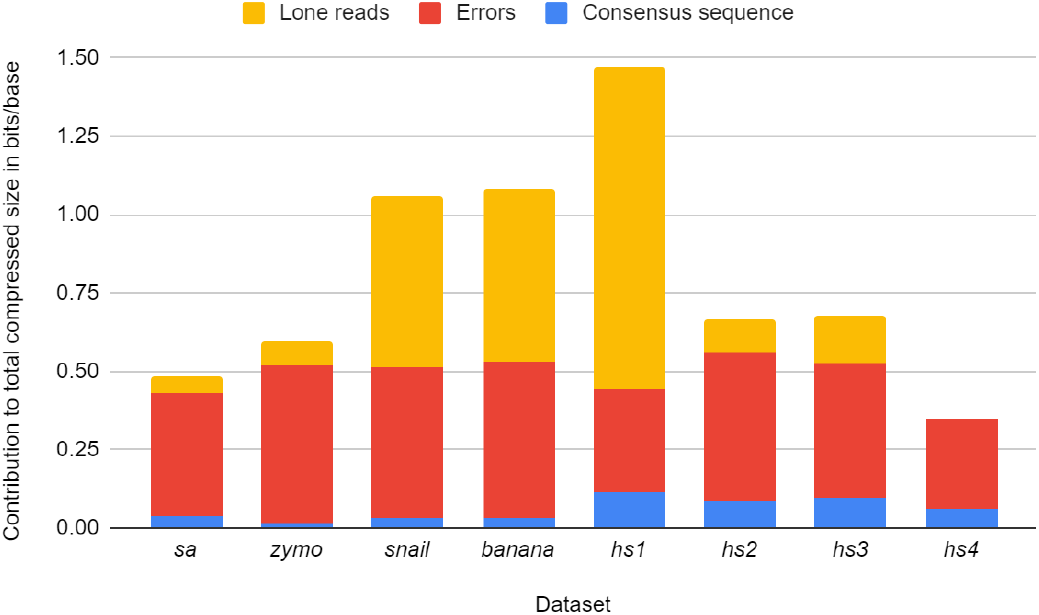
Contribution of consensus sequence, errors and lone reads to the compressed size across datasets.

#### Impact of varying coverage

To understand the impact of sequencing coverage on the performance of NanoSpring, we tested it on two datasets subsampled to multiple coverage values. We used the *banana* dataset with 177x coverage, and the CHM13 dataset (used to obtain *hs3* in Table 1) with 126x coverage. The compressed sizes are shown in Figure 3, where we see that the compression improves with coverage for both datasets even though the two datasets have significantly different compression levels (likely due to the differing basecalling qualities). This is expected since higher coverage datasets have more redundancy. For the CHM13 dataset, we get compressed sizes of 0.83, 0.68, 0.60, 0.57 and 0.56 bits/base for coverage values of 12x, 23x, 46x, 69x and 126x, respectively, with diminishing returns as the coverage increases. We observe that even at the low coverage of 12x, NanoSpring achieves more than two times better compression ratio than ENANO or Gzip for the high quality CHM13 dataset.

**Figure 3.**
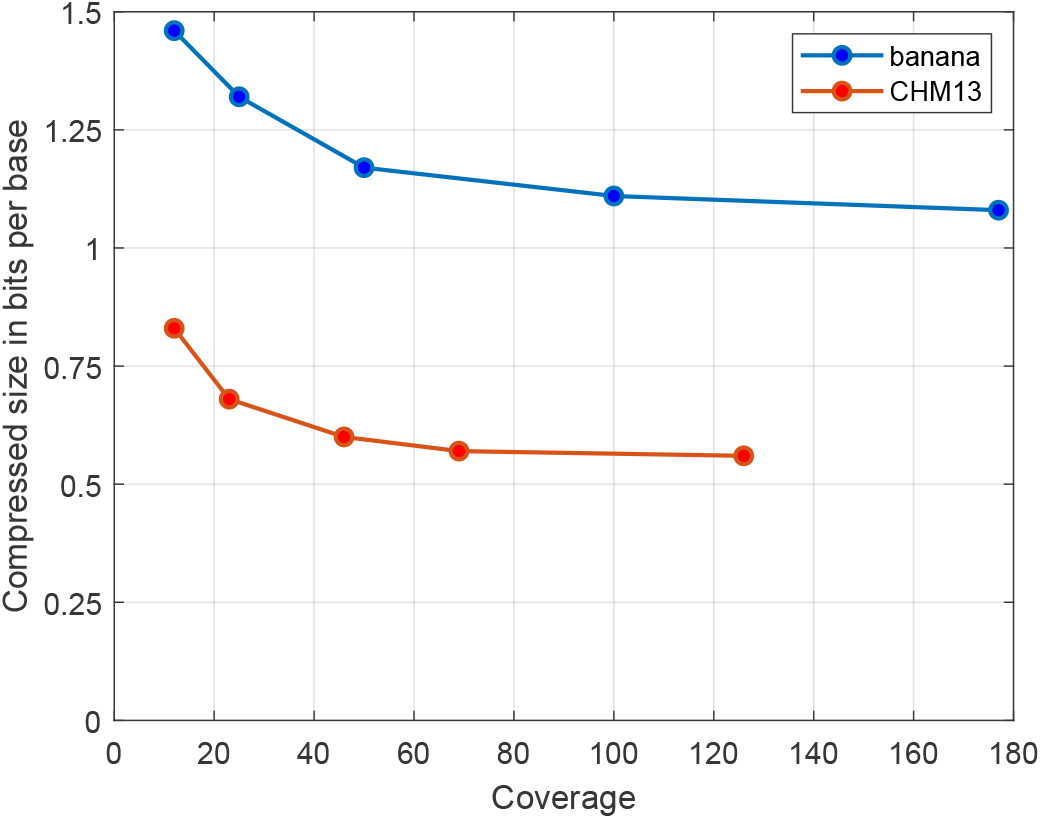
Compressed size (bits/base) vs. coverage for subsampled *banana* and CHM13 datasets.

#### Impact of basecalling error rate

To understand the impact of the basecaller error rate on the performance of NanoSpring, we basecalled the *S. aureus* dataset with three modes of the Guppy basecaller (version 6.1.2) by ONT: *fast, hac* (high-accuracy) and *sup* (super-accurate) (where we used the *hac* mode to obtain the *sa* dataset in Table 1). As shown in the Table 4, there is significant impact of the basecaller mode on the mean error rate and the compressed size. The compression time stays roughly constant while the memory usage increases for higher accuracies possibly because we get larger contigs in that case. For the *sup* mode with mean error rate below 4%, the compressed size is close to 0.4 bits/base, showing the potential for better compression as the basecaller quality continues to improve in the near future.

**Table 4.** Performance of NanoSpring for the *S. aureus* dataset under different basecaller modes.

| Basecaller mode | Mean error rate | Compressed size (bits/base) | Compression time | Peak memory usage (GB) |
| --- | --- | --- | --- | --- |
| *fast* | 7.01% | 0.695 | 28.7s | 5.3 |
| *hac* | 4.60% | 0.486 | 28.5s | 7.3 |
| *sup* | 3.68% | 0.410 | 27.7s | 7.2 |

## Methods

### MinHash indexing

We use MinHash^18^ for indexing the reads allowing for efficient lookup of reads overlapping with any given sequence. During the construction of the index, we first extract substrings of length *k* (k-mers) from a read and compute *n* pseudorandom hash functions of the k-mers (the MinHash *sketch* of the read). For each hash function, we find the k-mer with the minimum hash value, referred to as the MinHash of the read. The basic theoretical property underlying MinHash is that the fraction of shared MinHash values (out of *n*) between two sequences is a good estimator for the fraction of shared *k*-mers between the sequences, and the estimator accuracy increases with increasing *n*. Since we can expect overlapping sequences, potentially with substitution/insertion/deletion errors, to have common k-mers (for sufficiently small k), MinHash provides us a way to efficiently estimate the similarity of sequences and to rapidly look up overlapping reads (as described next).

The MinHash index consists of *n* tables, one for each pseudorandom hash function. Each table maps MinHash values for the corresponding hash function to the list of reads with that MinHash value. During lookup, we are given a sequence and first compute the *n* MinHash values for that sequence (the MinHash sketch). Then we use the index to find and return reads matching the sketch for at least *t* out of *n* hash functions, where *t* is a threshold parameter. In our implementation, the *k*-mers (with *k* ≤ 32) and hash values are represented as 64-bit integers, and the pseudorandom hash functions are simply computed as *hash*(*kmer* ⊕ *r_i_*) where *hash* is a standard hash function, and *r_i_* for *i* = 1,…, *n* are pseudorandom integers. To minimize the memory usage, the hash tables are built using BBHash^19^ which is a specialized data structure designed for hash tables that are not modified after construction, which applies in our case.

There are three key parameters for MinHash indexing, the *k*-mer length (*k*), the number of hash functions (*n*) and the threshold for lookup (*t*). Increasing *n* improves the accuracy of MinHash and allows for more reliable estimation of sequence similarity but leads to increased computational overhead. The ratio *t* /*n* determines the threshold for similarity of reads during lookup. Higher values of this ratio can lead to missed potentially matching reads, whereas smaller values can increase the number of false positives and hence the computational overhead. In addition, very small values can lead to low quality matches which can adversely affect the compression. Finally, the value of *k* should be chosen depending on the genome size, the error rate, and the computational requirements. At higher values of *k*, the *k*-mer is more likely to have an error and finding exact matches becomes unlikely. If *k* is too small, then we can get many spurious matches for large genomes leading to poor computational performance. By default, we set *k* = 23, *n* = 60, *t* = 6 (see Supplementary data for a detailed analysis). We note that the parameters chosen in this work are different from the parameters used in the MHAP assembler^13^. Specifically, we tuned the parameters for better performance since we only require an approximate assembly in our application. We also take into account the fact that the nanopore error rates have reduced significantly over the past years making the use of these parameters reasonable.

### Construction of contigs

NanoSpring uses the following pseudocode for constructing contigs of overlapping reads:

1. Initialization: Pick an arbitrary read not yet added to a contig, and construct the consensus graph with a single read.
2. Repeat the following until no matching reads are found:
  a. Obtain substring of the current consensus sequence (by shifting a average read-length sized window over the consensus sequence at each step, see below for a detailed explanation).
  b. Find candidate overlapping reads to the substring using MinHash index.
  c. For each candidate read from previous step (that has not yet been added to a contig):

i. Align read to current consensus sequence using minimap2.
ii. If alignment succeeds, add read to the consensus graph and recalculate the consensus sequence (explained below).

As described in the pseudocode, the contig construction maintains a consensus graph (also referred to as an assembly graph) and greedily adds reads to the graph. The graph is directed and acyclic with the nodes representing the bases and the edges storing the information about the reads passing through that edge. The *weight* of an edge is the number of reads passing through the edge. Initially the graph is just a line graph consisting of a single read. As reads get added to the graph, these reads lead to branching out from the line graph due to presence of errors. The consensus sequence represents the path with the highest weight (where the weight of a path is the sum of weights of the edges on the path). Figure 1a illustrates the consensus graph with the reads and the consensus sequence, and Figure 4 shows the overall contig generation procedure. In our implementation, we use a greedy algorithm for computing the consensus path: starting at the leftmost node and picking the highest weight edge at every step. We use the greedy algorithm instead of the optimal dynamic programming-based algorithm due to its simplicity and similar performance in practice.

**Figure 4.**
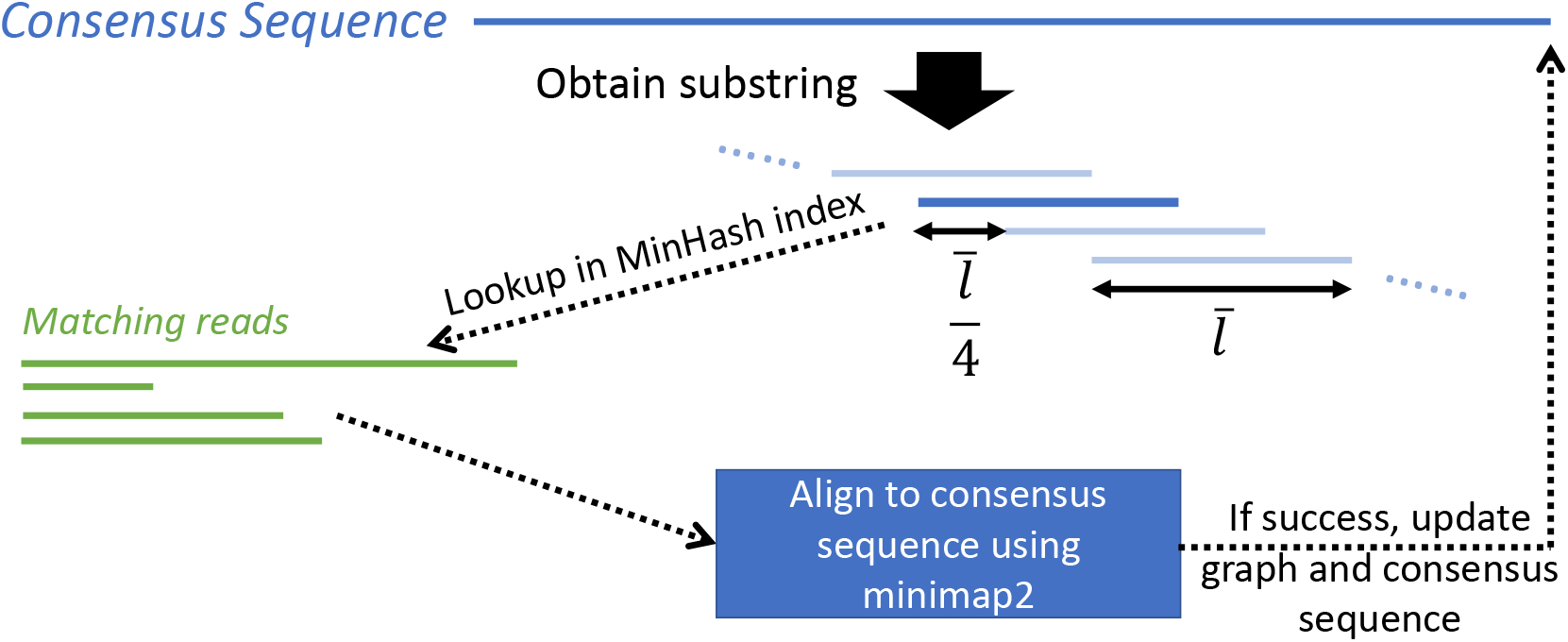
Contig generation process. A substring of the current consensus sequence is used to find matching reads from the MinHash index. Each potential match is aligned to the consensus sequence using minimap2 and added to the graph if the alignment succeeds. The substring has length 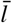 (average read length) and is shifted along the consensus sequence at each step to capture all potential matches.

At every step in the contig generation algorithm, we first pick a substring of the consensus sequence, which is used to search for matching reads using the MinHash index. The substring has length equal to the average read length of the dataset or the length of the consensus sequence, whichever is smaller. After each iteration, we obtain a shifted substring by changing the start position by a quarter of the average read length (shift length chosen based on experiments). We first shift the substring to the right until we reach the end, and then shift it left till we reach the beginning. This allows us to capture reads overlapping with any section of the consensus sequence. Note that the consensus sequence itself is constantly updated, and hence we maintain the position of the first read on the consensus sequence as a reference zero coordinate for tracking the substring location. We work with the consensus sequence at the current iteration (instead of an individual read) for MinHash lookup and alignment since we expect the consensus sequence to have a lower error rate leading to a more efficient and accurate process. To handle reverse complemented reads, we search for overlapping reads to both the substring and its reverse complement in step 2b and store a flag denoting reverse complemented reads. As a result, reads can pass through an edge in either direction.

For each potentially matching read obtained using MinHash, we attempt to align it to the consensus sequence using the minimap2 aligner^7^. We found that previous works on assembly^13^ used their own implementation of the optimal Myers aligner^20^, but we found that the widely used minimap2 aligner was simpler to use and significantly faster. The minimap2 aligner first indexes the reference sequence (the consensus sequence in our case) based on minimizers (lexicographically smallest hashed k-mers) of length *k* in each window of length w (both *k* and w are parameters). Then it attempts to locate these minimizers in the query string (the reads returned by MinHash), followed by more accurate alignment in the regions between these minimizers. When alignment succeeds, minimap2 returns the CIGAR string consisting of the errors in the query string with respect to the reference (we restrict ourselves to the top-scoring alignment returned by minimap2). We use this information to add the read to the graph, with soft clips treated as a sequence of insertions at the beginning or end of the read. We modified the default parameters for minimap2 in order to improve the computational performance, setting k = 20, w = 50 and reducing the `max-chain-iter` parameter controlling the complexity of the chaining step in minimap2 to 400 from the default value of 5000. See Supplementary data for more discussion on the impact of these parameters.

### Encoding and compression of streams

The reads in each contig are encoded into multiple streams after the contig generation is done. We store the consensus sequence of the contig (1 byte/base representation) and then store the representation of each read with respect to the consensus. Specifically, we store the start position of the read on the consensus sequence, the errors in the read, the reverse complement flag (bool), and the read index (32-bit unsigned integer) representing the index of the read in the original FASTQ file. The errors are themselves represented using three streams: (i) the error type (insertion/deletion/substitution) (char), (ii) the error position on the read (delta coded), and (iii) the erroneous base (for insertions and substitutions) (char). These streams are illustrated in Figure 1b. We observed that in many cases, we get multiple insertions at the beginning and end of the reads possibly due to adapter sequences (soft clipped in the alignment). For such cases, we simply encode the number of insertions (unsigned varint encoding) and the inserted bases (char) rather than encoding each insertion separately. The encoding of the insertions at the start and end of reads is done as part of the error position and erroneous base streams, where the error position stream for a read encodes (i) first the number of insertions at start, (ii) followed by the other error positions (delta coded), (iii) followed by the number of insertions at end. The current implementation combines the start position and error position streams since they share the same datatype (unsigned varint encoding) and the start position stream is typically a negligible contributor to the total size.

In our experiments, we found that the contig generation led to several contigs with only a single read due to the greedy procedure and the variation in read quality (with high error reads missed by MinHash). For any such contigs with just a single read, we directly write the read sequence using a text representation (1 byte/base) to a separate stream for these *lone* reads. These are not written to the consensus sequence stream.

Finally, these streams are compressed with general-purpose compressors and combined into a single file using the tar utility on UNIX. We use two general-purpose compressors which provide improvements over Gzip while being computationally efficient. For compressing the stream with the erroneous bases, we use LZMA2 (https://github.com/conor42/fast-lzma2) which relies on Lempel-Ziv compression^21^ and arithmetic coding^22^. For the remaining streams we use BSC (https://github.com/IlyaGrebnov/libbsc) which relies on the Burrows-Wheeler transform (BWT^23^) and arithmetic coding. As discussed in Supplementary data, we found this combination of compressors led to the best compression ratios.

### Additional implementation details

We made some modifications to the procedure presented above to improve the performance on real datasets.

- The MinHash indexing, contig generation, stream compression, and the decompression are parallelized to improve the wall-clock performance. During contig generation, different threads work on different contigs, and we ensure that there are no conflicts using locks. The impact of multithreading is discussed in Supplementary data.
- We found that for certain human datasets, minimap2 took a very long time for aligning highly repetitive sequences, usually with tandem repeats (such as *GT GT GT…*). Therefore, we check the reads for short tandem repeats before the contig generation stage and repetitive reads are directly written to the lone read stream. We also write reads with length ≤ 32 directly to the lone read stream since they have too few k-mers to obtain matches using MinHash.
- During compression, we store the reads encoded using 2 bits/base on disk to limit the memory usage. We found that the disk based random access to load these reads for minimap2 alignment is sufficiently fast to justify the reduction in memory usage (around 22 GB memory reduction for a 30x human dataset).
- To limit the memory usage of the consensus graphs during contig generation, especially when working with multiple threads, we impose a limit on the number of edges in the graph. Beyond this limit, the contig generation is stopped and we proceed with a new contig. We found that this simple strategy drastically reduces the peak memory consumption while having minimal impact on the compression ratio (see Supplementary data). We also use other low-level optimizations such as periodically calling `malloc_trim()` to further reduce the memory usage.
- During compression the reads are encoded in the order they appear in the contigs. Thus the decompressor needs to reorder these reads using the stored index to recover the original order of reads in the FASTQ. We use efficient disk-based sorting to limit the memory usage of decompression to around 5 GB irrespective of the file size.

## Discussion

We presented NanoSpring, a reference-free compressor for nanopore sequencing reads, relying on an approximate assembly approach. We evaluate NanoSpring on a variety of datasets including bacterial, metagenomic, plant, animal, and human whole genome data. For recently basecalled high quality nanopore datasets, NanoSpring, which focuses only on the base sequences in the FASTQ file, uses just 0.35-0.7 bits per base which is 3-6x lower than general purpose compressors like gzip. NanoSpring is competitive in compression ratio and compression resource usage with the state-of-the-art tool CoLoRd while being significantly faster at decompression. NanoSpring is open-source and available on GitHub at https://github.com/qm2/NanoSpring. Future work includes research on the assembly algorithm to get closer to the fundamental limits. Another important direction is to incorporate NanoSpring into a full-fledged FASTQ compressor capable of handling quality scores and read identifiers, possibly by combining the best aspects of ENANO and NanoSpring.

## Acknowledgements

We thank the Stanford EE REU program for providing the opportunity to initiate this project and acknowledge funding from Philips.

## Author contributions statement

T.W. and S.C. conceived of the idea. Q.M., S.C. and Y.Z. were involved in designing, implementing and testing the proposed algorithm. Q.M. ran the experiments for the manuscript. Q.M. and S.C. prepared the draft manuscript. All authors read and approved the final manuscript.

## Data and code availability statement

All the datasets used in the experiment are publicly available and instructions for accessing the same are provided in the supplementary material. In addition, all the tools used for the experiments, including NanoSpring, are open source and can be obtained as described in the supplementary material.

## Additional information

The author(s) declare no competing interests.

## Notes

https://github.com/qm2/NanoSpring

### Competing Interest Statement

The authors have declared no competing interest.

### Summary of Updates

- update NanoSpring to use less memory, other bug fixes - compare with CoLoRd compressor - added new datasets - reformatting

