## Supplementary Data for "NanoSpring: reference-free lossless compression of nanopore sequencing reads using an approximate assembly approach"

GitHub: `https://github.com/qm2/NanoSpring`

### 1 Datasets

In this section, we provide instructions for obtaining the datasets used for evaluating NanoSpring and the other tools. While NanoSpring supports gzipped input FASTQ during compression, we decompressed to the FASTQ file for a fair comparison with the other tools.

#### 1.1 *S. aureus*

The *S. aureus* dataset was obtained from the work in Wick *et al.* (2019) where it was used for evaluating basecalling tools. We downloaded the fast5 dataset with the raw signal and basecalled it with three basecaller modes for Guppy 6.1.2. We also calculated the mean error rate using alignment. Note that we use the basecall with the *hac* mode as the dataset for the main results (*sa* dataset).

##### Downloading the fast5 files and the reference sequence

```
mkdir Staphylococcus_aureus_CAS38_02/
cd Staphylococcus_aureus_CAS38_02/
mkdir fast5/ && cd fast5/
wget -O Staphylococcus_aureus_CAS38_02_fast5s.tar.gz \
https://bridges.monash.edu/ndownloader/files/14260568
tar -xzvf Staphylococcus_aureus_CAS38_02_fast5s.tar.gz
cd ../
wget -O Staphylococcus_aureus_CAS38_02_reference.fasta.gz \
https://bridges.monash.edu/ndownloader/files/14260241
gunzip Staphylococcus_aureus_CAS38_02_reference.fasta.gz
rm fast5/Staphylococcus_aureus_CAS38_02_fast5s.tar.gz
cd ../
```

##### Basecalling

```
wget https://cdn.oxfordnanoportal.com/software/analysis/ont-guppy_6.1.2_linux64.tar.gz
tar -xzvf ont-guppy_6.1.2_linux64.tar.gz
rm ont-guppy_6.1.2_linux64.tar.gz
for config in "fast" "hac" "sup"; do
    ./ont-guppy/bin/guppy_basecaller -i Staphylococcus_aureus_CAS38_02/fast5/ \
    -s tmp_out_$config/ -c dna_r9.4.1_450bps_$config.cfg -x cuda:all
    cat tmp_out_$config/pass/*.fastq tmp_out_$config/fail/*.fastq >\
    Staphylococcus_aureus_CAS38_02/basecall_$config.fastq
    rm -r tmp_out_$config/
done
```

##### Alignment and error calculation

We use minimap2 (Li, 2018) and Samtools (`http://www.htslib.org/`) for calculating the mean error rates for the three basecaller modes.

```
for config in "fast" "hac" "sup"; do
    minimap2 -ax map-ont \
    Staphylococcus_aureus_CAS38_02/Staphylococcus_aureus_CAS38_02_reference.fasta \
    Staphylococcus_aureus_CAS38_02/basecall_$config.fastq -t 20 | samtools stats - > \
    Staphylococcus_aureus_CAS38_02/basecall_$config.stats.txt
done
```

We use the `error rate` entry in the generated statistics file as the mean error rate.

#### 1.2 Zymo

The Zymo dataset (Nicholls *et al.*, 2019) is a metagenomic dataset and we use the bonito basecalled version as used in Kokot *et al.* (2022).

```
wget ftp://ftp.sra.ebi.ac.uk/vol1/fastq/ERR539/000/ERR5396170/ERR5396170.fastq.gz
```

#### 1.3 Snail

The snail dataset (Sun *et al.*, 2021) was obtained from `https://www.ebi.ac.uk/ena/browser/view/SRR12763791`.

```
wget ftp://ftp.sra.ebi.ac.uk/vol1/fastq/SRR127/091/SRR12763791/SRR12763791_1.fastq.gz
```

According to the reference, the dataset was basecalled using Guppy 3.6.0 (high accuracy mode).

#### 1.4 Banana

The banana dataset (Belser *et al.*, 2021) was obtained from `https://www.ebi.ac.uk/ena/browser/view/ERX5238183`.

```
wget ftp://ftp.sra.ebi.ac.uk/vol1/fastq/ERR545/008/ERR5455028/ERR5455028.fastq.gz
```

According to the reference, the dataset was basecalled using Guppy 4.0.1.

The original 177x dataset was used as the *banana* dataset in the main results. In addition we obtained several subsampled versions for studying the impact of coverage on the performance of NanoSpring. We used seqtk (`https://github.com/lh3/seqtk`) for generating the subsampled dataset. The command used is

```
seqtk sample -s100 ERR5455028.fastq.gz $fraction_reads > ERR5455028_$fraction_reads.fastq
```

where the variable `fraction_reads` was set to 0.0705, 0.141, 0.282 and 0.565 to get subsampled datasets of coverage approximately 12x, 25x, 50x and 100x, respectively.

#### 1.5 Human: NA12878

The human NA12878 dataset (Jain *et al.*, 2018) was obtained from `https://github.com/nanopore-wgs-consortium/NA12878/blob/master/Genome.md`, where we use the latest rel6 version basecalled with Guppy 2.3.8.

```
wget http://s3.amazonaws.com/nanopore-human-wgs/rel6/rel_6.fastq.gz
```

#### 1.6 Human: GM24385 (*hs2*)

The human GM24385 dataset (*hs2*) was obtained from `https://labs.epi2me.io/gm24385_2020.09/` and is part of a series of high quality open source datasets released by Oxford Nanopore Technologies (ONT) at `https://labs.epi2me.io/dataindex`. We use part of the dataset from September 2020 basecalled using Guppy 4.0.11 (high accuracy mode), downloaded using the following command.

```
aws s3 cp --no-sign-request \
s3://ont-open-data/gm24385_2020.09/analysis/r9.4.1/\
20200914_1357_1-E11-H11_PAF27462_d3c9678e/guppy_v4.0.11_r9.4.1_hac_prom/basecalls.fastq.gz\
 .
```

#### 1.7 Human: CHM13

The human CHM13 dataset (Nurk *et al.*, 2021) is part of the telomere-to-telomere (T2T) consortium attempting to sequence and assemble the human genome without any gaps. The dataset was obtained from `https://github.com/marbl/CHM13`, where we use the latest rel7 version basecalled with Bonito 0.3.1.

```
wget https://s3-us-west-2.amazonaws.com/human-pangenomics/T2T/CHM13/nanopore/rel7/rel7.fastq.gz
```

The original 126x dataset was used to obtain several subsampled versions for studying the impact of coverage on the performance of NanoSpring. We used seqtk (`https://github.com/lh3/seqtk`) for generating the subsampled dataset. The command used is

```
seqtk sample -s$seed rel7.fastq.gz $fraction_reads > rel7_$fraction_reads.fastq
```

where seed was set to 100 and variable `fraction_reads` was set to 0.099, 0.198 and 0.595 to get subsampled datasets of coverage approximately 12x, 23x and 69x, respectively. The seed was set to 11 and variable was set to 0.397 to get subsampled dataset of coverage approximately 46x. The 23x dataset corresponds to the *hs3* in the main result.

##### 1.8 Human: GM24385 (*hs4*)

The human GM24385 dataset (*hs4*) was obtained from `https://labs.epi2me.io/gm24385_q20_2021.10/` and is part of a series of high quality open source datasets released by Oxford Nanopore Technologies (ONT) at `https://labs.epi2me.io/dataindex`. We use part of the R10.4 dataset from October 2021 basecalled using Guppy 5.0.15 (super accuracy mode), downloaded using the following command (the two files were concatenated for our experiments).

```
aws s3 cp --no-sign-request \
s3://ont-open-data/gm24385_q20_2021.10/analysis/20210805_1713_5C_PAH79257_0e41e938/\
guppy_5.0.15_sup/basecalls.fastq.gz part_1.fastq.gz

aws s3 cp --no-sign-request \
s3://ont-open-data/gm24385_q20_2021.10/analysis/20210805_1713_5D_PAH78823_61ef97d4/\
guppy_5.0.15_sup/basecalls.fastq.gz part_2.fastq.gz
```

##### 1.9 N50 calculation

The script for calculating the N50 read length for a FASTQ file is available at `https://github.com/qm2/NanoSpring/util/sctipts/get_fastq_stats.sh`.
The command used to calculate the N50 for `file.fastq` is

```
cat file.fastq | sed -n '2~4p' | awk '{ print length }' | ./get_fastq_stats.sh
```

The command used to calculate the N50 for `file.fastq.gz` (gzipped FASTQ file) is

```
zcat file.fastq.gz | sed -n '2~4p' | awk '{ print length }' | ./get_fastq_stats.sh
```

#### 2 Installing and running tools

In this section, we provide instructions for running the various tools used in the experiments. The scripts used for running the main experiments and the parameter evaluation are available at `https://github.com/qm2/NanoSpring/tree/master/util/scripts`. The logs for the experiments are available at `https://github.com/qm2/NanoSpring/tree/master/logs`. Further information on installing the tools in various environments can be found on their respective GitHub pages.

##### 2.1 NanoSpring

###### 2.1.1 Download and install

Commit:
`https://github.com/qm2/NanoSpring/commit/919e5e13a077a6185997b73a87ff0509944c4581`

```
git clone --recursive https://github.com/qm2/NanoSpring.git
cd NanoSpring
mkdir build
cd build
cmake ..
make -j
```

###### 2.1.2 Usage

General usage:

```
Allowed options:
  -h [ --help ]                     produce help message
  -c [ --compress ]                 compress
  -d [ --decompress ]               decompress
  -i [ --input-file ] arg           input file name
  -o [ --output-file ] arg          output file name
  -t [ --num-threads ] arg (=20)    number of threads (default 20)
  -k [ --kmer ] arg (=23)           kmer size for the minhash (default 23)
  -n [ --num-hash ] arg (=60)       number of hash functions for minhash (default
                                    60)
  --overlap-sketch-thr arg (=6)     the overlap sketch threshold for minhash
                                    (default 6)
  --minimap-k arg (=20)             kmer size for the minimap2 (default 20)
  --minimap-w arg (=50)             window size for the minimap2 (default 50)
  --max-chain-iter arg (=400)       the max number of partial chains during
                                    chaining for minimap2 (default 400)
  --edge-thr arg (=4000000)         the max number of edges allowed in a consensus
                                    graph (default 4000000)
  -w [ --working-dir ] arg (=.)     directory to create temporary files (default
                                    current directory)
  --decompression-memory arg (=5)   attempt to set peak memory usage for
                                    decompression in GB (default 5 GB) by using
                                    disk-based sort for writing reads in the
                                    correct order. This is only approximate and
                                    might have no effect at very low settings or
                                    with large number of threads when another
                                    decompressor stage is the biggest memory
                                    contributor. Very low values might lead to
                                    slight reduction in speed.
```

For compressing `file.fastq` using default 20 threads (used for the main experiments):

```
./NanoSpring -c -i file.fastq -o file.NanoSpring
```

For compressing `file.fastq.gz` (gzipped fastq file) using default 20 threads:

```
./NanoSpring -c -i file.fastq.gz -o file.NanoSpring
```

Decompressing the `file.NanoSpring` with default 20 threads to `file.reads` (used for the main experiments):

```
./NanoSpring -d -i file.NanoSpring -o file.reads
```

More examples are provided on the GitHub README.

#### 2.2 pigz

##### 2.2.1 Download and install

```
wget https://zlib.net/pigz/pigz-2.4.tar.gz
tar -xzvf pigz-2.4.tar.gz
cd pigz-2.4
make
```

##### 2.2.2 Usage

Compression with 20 threads:

```
./pigz-2.4/pigz -k -p 20 file.fastq
```

Decompression with 20 threads:

```
./pigz-2.4/unpigz -k -p 20 file.fastq.gz
```

#### 2.3 ENANO

##### 2.3.1 Download and install

Commit:
https://github.com/guilledufort/EnanoFASTQ/commit/d6119dbccd19485c8222ec6d72516114401d1a08

```
git clone https://github.com/guilledufort/EnanoFASTQ.git
cd EnanoFASTQ/enano
make
```

##### 2.3.2 Usage

Compression with 20 threads:

```
./enano -t 20 file.fastq file.enano
```

Decompression with 20 threads:

```
./enano -d -t 20 file.enano file_decompressed.fastq
```

#### 2.4 CoLoRd

##### 2.4.1 Download and install

Commit:
https://github.com/refresh-bio/colord/commit/91d70e9d74a9138c4f2a9567eed82fa4899f6713

```
git clone https://github.com/refresh-bio/colord
cd colord && make
cd bin
```

##### 2.4.2 Usage

Compression with 20 threads in the balanced mode with quality and read identifier compression mode set to none.:

```
./colord compress-ont -p balanced -t 20 -q none -i none file.fastq file.colord
```

Decompression:

```
./colord decompress file.colord file_decompressed.fastq
```

#### 3 Parameter Experiment

To provide further insight into the NanoSpring algorithm, we discuss various parameters, experiments and related observations. While testing the individual parameters, we keep all other parameters constant. We note that the time measurements show some experimental variation due to I/O and multithreading. Therefore, we focus on the overall trend rather than these minor variations. More details on these experiments are available on GitHub.

##### 3.1 MinHash parameters

| $k$ | Compressed size | | Compression time | | Peak memory usage | |
|---|---|---|---|---|---|---|
| | *hs2* | *zymo* | *hs2* | *zymo* | *hs2* | *zymo* |
| 10 | 1.558 | 0.597 | 7h19m | 1h39m | 22.0 | 17.3 |
| 12 | 0.851 | 0.597 | 4h20m | 1h32m | 18.6 | 17.6 |
| 15 | 0.689 | 0.595 | 3h55m | 1h30m | 17.7 | 18.6 |
| 18 | 0.646 | 0.598 | 3h37m | 1h33m | 17.9 | 18.5 |
| 20 | 0.655 | 0.597 | 3h24m | 1h35m | 18.0 | 18.5 |
| 23 | 0.668 | 0.602 | 3h16m | 1h21m | 18.1 | 18.5 |
| 25 | 0.665 | 0.603 | 3h11m | 1h42m | 18.4 | 18.3 |
| 30 | 0.694 | 0.607 | 2h45m | 1h54m | 18.4 | 19.0 |

Table 1: Performance of NanoSpring with different MinHash $k$-mer sizes for the *hs*2 and *zymo* datasets. Compressed size is in bits/base and the peak memory usage is in GB.

| $n$ | $t$ | $t/n$ | Compressed size | | Compression time | | Peak memory usage | |
|---|---|---|---|---|---|---|---|---|
| | | | *hs2* | *zymo* | *hs2* | *zymo* | *hs2* | *zymo* |
| 50 | 4 | 0.08 | 0.661 | 0.605 | 3h15m | 1h43m | 20.2 | 18.6 |
| 50 | 5 | 0.10 | 0.658 | 0.600 | 3h05m | 1h23m | 18.6 | 18.3 |
| 60 | 6 | 0.10 | 0.656 | 0.601 | 3h08m | 1h22m | 19.0 | 18.6 |
| 70 | 7 | 0.10 | 0.654 | 0.600 | 2h59m | 1h23m | 18.9 | 19.6 |
| 50 | 6 | 0.12 | 0.687 | 0.599 | 2h52m | 1h19m | 17.7 | 19.3 |

Table 2: Performance of NanoSpring with varying MinHash $t$ and $n$ parameters for the *hs*2 and *zymo* datasets. Compressed size is in bits/base and the peak memory usage is in GB.

NanoSpring uses MinHash to index the reads and find overlapping reads during contig generation, with parameters $k$ (MinHash $k$-mer length), $n$ (number of hash functions) and $t$ (threshold number of matches for successful lookup). In Table 1, we look at the performance for the *hs2* and *zymo* datasets with varying $k$ (default is $k = 23$). The impact of $k$ is dependent on the genome size since for a larger genome we expect to get more false positive matches as $k$ decreases. For the *hs2* dataset, we see that the compression time increases as $k$ decreases. The compression first improves with decreasing $k$ since smaller $k$-mers are less likely to have errors leading to fewer missed matches. But as $k$ decreases further, the compression worsens due to increasing false positive matches. For the *zymo* dataset which consists of bacterial and yeast genomes, we see that the compression improves until $k = 20$. Thus, the value of $k$ should be chosen based on the genome size and error rate, with the default value of $k = 23$ chosen to give a reasonable tradeoff for most human datasets.

Table 2 shows the performance for the same two datasets as the parameters $n$ and $t$ are varied (default values $n = 60, t = 6$). As the ratio $t/n$ increases, we miss more and more matches, while very small values lead to false positive matches and increased computation time for minimap2 alignment. While the best value of this ratio depends on the dataset and the error rate, we found the ratio of 0.10 to work consistently well. For a fixed ratio $t/n$, we expect higher $n$ to give better results (Berlin *et al.*, 2015) at the cost of more computation. In practice we found that the dependence of compression ratio on $n$ was not significant within a range of values, and we chose $n = 60$ based on experiments across datasets. We also note some variation in memory usage with the three MinHash parameters, although this is less pronounced that the variation in time and compression ratio.

#### 3.2 Minimap2 parameters

| $k$ | $w$ | $mci$ | Compressed size | | Compression time | | Peak memory usage | |
|---|---|---|---|---|---|---|---|---|
| | | | *hs2* | *hs3* | *hs2* | *hs3* | *hs2* | *hs3* |
| 15 | 10 | 5000 | 0.678 | 0.709 | 7h43m | 16h00m | 18.5 | 25.4 |
| 15 | 10 | 400 | 0.669 | 0.706 | 5h12m | 6h40m | 19.2 | 20.2 |
| 20 | 50 | 5000 | 0.653 | 0.682 | 3h30m | 4h55m | 18.6 | 19.4 |
| 20 | 50 | 400 | 0.655 | 0.681 | 3h05m | 3h48m | 18.6 | 21.6 |

Table 3: Performance of NanoSpring with varying minimap2 $k$, $w$ and $mci$ (`max-chain-iter`) parameters for the $hs2$ and $hs3$ datasets. Compressed size is in bits/base and the peak memory usage is in GB.

NanoSpring uses the minimap2 aligner (Li, 2018) to align candidate reads to the consensus sequence and add them to the graph during contig generation. We tuned the default minimap2 parameters to improve the performance of the algorithm, focusing on three main parameters: the minimizer $k$-mer size, the minimizer window size $w$ and the maximum number of iterations during the chaining step (`max-chain-iter` or $mci$). We changed the default minimap2 setting of $(k = 15, w = 10, mci = 5000)$ to $(k = 20, w = 50, mci = 400)$ in order to get the best tradeoff between compressed size and compression time. Table 3 shows the results for four parameter settings for the *hs2* and *hs3* datasets. We observe that the parameters have a significant effect on compression time, while the impact on compressed size is much smaller. The choice of the appropriate parameters was especially crucial for the *hs3* dataset, which requires 4 times more time with the default minimap2 setting as compared to the setting used in NanoSpring (the memory usage also improves for the chosen parameters). Based on some tests, we believe that this effect is due to the presence of more repetitive sequences in the *hs3* dataset which leads to extremely slow alignment unless the `max-chain-iter` parameter is reduced.

#### 3.3 Stream compressor

| Compressor | Compressed size (bits/base) |
|---|---|
| Gzip | 0.878 |
| BSC | 0.696 |
| LZMA2 | 0.689 |
| BSC+LZMA2 | 0.668 |

Table 4: Effect of stream compressor on compressed size for the $hs2$ dataset. In the last row which corresponds to the default setting for NanoSpring, the stream of erroneous bases is compressed using LZMA2 and the other streams are compressed using BSC.

Table 4 shows the effect of the stream compressor on the overall compression ratio. NanoSpring uses LZMA2 for compressing the stream of erroneous bases and BSC for the other streams. We see that Gzip performs around 30% worse than the default setting. But NanoSpring with Gzip still has 2x better compression than ENANO suggesting that the advantage of NanoSpring is mostly due to the approximate assembly process. Between BSC and LZMA2, we found that BSC was better for most streams, but LZMA2 was around 10% better for the erroneous base stream which is a major contributor to the overall compressed size for certain datasets. Thus, we use a combination of the two for best results. We note that the computational requirements for stream compression are relatively small as compared to the approximate assembly process and were not a major factor in the choice of the compressor. The only exception to this was the memory usage during decompression, where choosing a reasonably small block for BSC is important (we use 48 MB block size).

#### 3.4 Threads

| Number of threads | Compressed size (bits/base) | Compression time | Peak memory usage (GB) |
|---|---|---|---|
| 5 | 0.671 | 8h57m | 7.89 |
| 10 | 0.665 | 5h00m | 12.1 |
| 20 | 0.663 | 3h09m | 18.7 |
| 30 | 0.654 | 2h37m | 26.3 |

Table 5: Performance of NanoSpring with different number of threads for the *hs*2 dataset.

Table 5 shows the compression performance of NanoSpring with different number of threads on the *hs2* dataset (where we used 20 threads by default for the main experiments). The time required for compression reduces as we use more threads, while the peak memory usage increases because the threads work simultaneously on different contigs. We observe that the compressed size is roughly constant with the increasing thread count. There is a slight impact of multithreading because the threads compete for reads to be added in their respective contigs, and more threads means that the contigs are more fragmented. For even higher number of threads, we expect the disk I/O and memory allocations to become the bottleneck leading to diminishing returns.

#### 3.5 Edge threshold

| Edge threshold | Compressed size (bits/base) | Compression time | Peak memory usage (GB) |
|---|---|---|---|
| 1M | 0.699 | 2h50m | 8.9 |
| 4M | 0.662 | 3h07m | 18.7 |
| 16M | 0.669 | 3h49m | 52.0 |
| $\infty$ | 0.652 | 4h10m | 100.4 |

Table 6: Performance of NanoSpring with different edge thresholds for the *hs*2 dataset.

To reduce the peak memory usage during compression, NanoSpring imposes a limit on the number of edges in a consensus graph during contig generation (4 million by default). Table 6 shows the compression results for the *hs2* dataset for different values of the threshold. The last row shows the result without any edge threshold, and we see the memory usage is more than 100 GB as compared to roughly 20 GB for the default threshold. In general, we see that decreasing the threshold significantly reduces the memory usage and slightly reduces the time but leads to more fragmented contigs and worse compression. We selected 4M as the default threshold to achieve a practical tradeoff between these factors.
